# Polymorphic positions 349 and 725 of the autoimmunity-protective allotype 10 of ER aminopeptidase 1 are key in determining its unique enzymatic properties

**DOI:** 10.1101/2024.03.14.584938

**Authors:** Galateia Georgaki, Anastasia Mpakali, Myrto Trakada, Athanasios Papakyriakou, Efstratios Stratikos

## Abstract

ER aminopeptidase 1 (ERAP1) is a polymorphic intracellular aminopeptidase with key roles in antigen presentation and adaptive immune responses. ERAP1 allotype 10 is highly protective towards developing some forms of autoimmunity and displays unusual functional properties, including very low activity versus some substrates. To understand the molecular mechanisms that underlie the biology of allotype 10 we studied its enzymatic and biophysical properties focusing on its unique polymorphisms V349M and Q725R. Compared to ancestral allotype 1, allotype 10 is much less effective in trimming small substrates but presents allosteric kinetics that ameliorate activity differences at high substrate concentrations. Furthermore, it is inhibited by a transition-state analogue via a non-competitive mechanism and is much less responsive to an allosteric small-molecule modulator. It also presents opposite enthalpy, entropy and heat capacity of activation compared to allotype 1 and its catalytic rate is highly dependent on viscosity. Polymorphisms V349M and Q725R significantly contribute to the lower enzymatic activity of allotype 10 for small substrates, especially at high substrate concentrations, influence the cooperation between the regulatory and active sites and regulate viscosity dependence, likely by limiting product release. Overall, our results suggest that allotype 10 is not just an inactive variant of ERAP1 but rather carries distinct enzymatic properties that largely stem from changes at positions 349 and 725. These changes affect kinetic and thermodynamic parameters that likely control rate-limiting steps in the catalytic cycle, resulting in an enzyme optimized for sparing small substrates and contributing to the homeostasis of antigenic epitopes in the ER.

## INTRODUCTION

Polymorphic variation in immune system genes underlies the variability of immune responses in natural populations and predisposition to infections, cancer, and autoimmunity. Amongst the most genetically variable biochemical pathways of the immune system is the pathway of antigen processing and presentation. The Major Histocompatibility Class I molecules (MHC-I) are highly polymorphic with tens of thousands of haplotypes discovered to date^1^ which influence the binding of small peptides derived from the degradation of intracellular or endocytosed extracellular proteins. The complexes of MHC-I with these peptides are displayed on the cell surface of all somatic cells and constitute a snapshot of the cell proteome^2^. Changes in displayed peptides may indicate infection or cellular transformation and are recognized by specific receptors on cytotoxic T-cells initiating cascades that lead to the death of the recognized cell. Intracellular aminopeptidases, such as Endoplasmic Reticulum Aminopeptidase 1 (ERAP1), prepare the peptidic cargo of MHC-I by trimming the N-terminus of precursor peptides down to the optimal length or over-trimming mature peptides limiting their opportunity to bind onto MHC-I^3^.

ERAP1 is primarily found in the ER of most somatic cells and is an integral part of the antigen processing and presentation pathway^3^. Its activity can regulate the repertoire of peptides that are bound and presented by MHC-I molecules (collectively referred to as the immunopeptidome) and can thus regulate adaptive immune responses in both health and diseases. ERAP1 function has been linked to the predisposition to several autoinflammatory diseases of autoimmune aetiology (in particular MHC-I-opathies^4^), the efficacy of immune responses to several infections^5,6^ as well as to the efficacy of cancer immunotherapy^7^, and as a result it is an emerging pharmacological target^8,9^. ERAP1 is also polymorphic and several missense coding single nucleotide polymorphisms (SNPs) have been associated with predisposition to disease^10^ while inducing significant changes in enzymatic function^11,12^. The nine most common ERAP1 SNPs are, however, not distributed randomly in the human population but rather form a few common allotypes^13,14^. These ERAP1 allotypes present significant functional differences that may underlie their association with disease predisposition^14^. Of the ten most common ERAP1 allotypes, allotype 10 (often referred to as Hap10) is a strong functional outlier that is found in approximately 22% of humans and has been reported to be protective for some forms of autoimmunity such as Psoriasis and Behçet’s^15,16^. ERAP1 allotype 10 has been shown to have much lower enzymatic activity (up to 60-fold) when measured using some substrates^14^. This has led several researchers to refer to ERAP1 allotype 10 as a sub-active or even an inactive ERAP1 variant^16–18^. However, a recent study by our group following the parallel trimming of hundreds of peptides *in vitro* suggested that Hap10 is not inactive but rather has distinct substrate specificity^19^. The molecular basis of the unique enzymatic properties of this important ERAP1 allotype is currently poorly understood. Still, inspection of the amino acid composition of common ERAP1 allotypes indicates that allotype 10 consists of two SNPs that are unique only to this allotype, namely V349 and Q725 (Table 1).

**Table 1:** SNP composition of common ERAP1 allotypes. Data taken from ^14^. Polymorphic positions unique to allotype 10 are highlighted.

| Amino acid composition at each SNP |  |  |  |  |  |  |  |  |  |
| --- | --- | --- | --- | --- | --- | --- | --- | --- | --- |
| Allotype | 56 | 127 | 276 | 346 | 349 | 528 | 575 | 725 | 730 |
| *1(ancestral)* | E | P | I | G | M | K | D | R | Q |
| 2 | E | R | I | G | M | K | D | R | Q |
| 3 | E | R | I | G | M | K | D | R | E |
| 4 | E | R | I | G | M | R | D | R | E |
| 5 | E | R | I | D | M | R | D | R | E |
| 6 | E | P | I | G | M | R | D | R | E |
| 7 | K | P | I | G | M | R | D | R | E |
| 8 | E | P | M | G | M | R | D | R | E |
| 9 | E | P | M | G | M | R | N | R | E |
| **10** | E | P | I | G | **V** | R | N | **Q** | E |

In this study, we used kinetic and biophysical analyses to investigate the properties of allotype 10 in comparison to the ancestral allotype 1. We focused our comparison on positions 349 and 725 which are unique to allotype 10 under the hypothesis that these SNPs may be critical in shaping the properties of this allotype. Our results suggest that allotype 10 displays distinct kinetic and thermodynamic properties including allosteric behaviour and an altered connection between the regulatory and active sites. Positions 349 and 725 are important for determining the activity range of allotype 10 but do not determine its allosteric kinetic behaviour. Overall, our data indicate that allotype 10 is an ERAP1 variant optimized to trim larger peptides while sparing smaller ones, which would place this allotype in a unique position to influence both antigen presentation and peptide homeostasis in the ER.

## MATIERALS AND METHODS

### Plasmids

The coding sequence for ERAP1 allotype 1 and allotype 10 containing a C-terminal 10-His purification tag next to a TEV recognition site was obtained by custom gene synthesis (BioCat GmbH, Heidelberg). The final construct has a length of 960 amino acids and a MW of 109585 kDa (allotype 1) or 109553 kDa (allotype 10) . The codon-optimised ERAP1 gene was synthesized and sub-cloned into a pFastBac1 vector using BamH1 and XhoI to generate pFASTBAC1-ERAP1_Hap1 and pFASTBAC1-ERAP1_Hap10 as shown below (signal sequence is underlined; TEV recognition site is underlined in bold font and His tag in italics; common SNPs in bold).

pFASTBAC1-ERAP1_Hap1: <u>MVFLPLKWSLAIMSFLLSSLLALLTVSTPSWC</u>QSTEASPKRSDGTPFPWNKIRLP**E**YVIPVH YDLLIHANLTTLTFWGTTKVEITASQPTSTIILHSHHLQISRATLRKGAGERLSEEPLQVLE HP**P**QEQIALLAPEPLLVGLPYTVVIHYAGNLSETFHGFYKSTYRTKEGELRILASTQFEPTA ARMAFPCFDEPAFKASFSIKIRREPRHLAISNMPLVKSVTVAEGLIEDHFDVTVKMSTYLVA FIISDFESVSKITKSGVKVSVYAVPDK**I**NQADYALDAAVTLLEFYEDYFSIPYPLPKQDLAA IPDFQSGAMENWGLTTYRESALLFDAEKSSASSKL**G**IT**M**TVAHELAHQWFGNLVTMEWWNDL WLNEGFAKFMEFVSVSVTHPELKVGDYFFGKCFDAMEVDALNSSHPVSTPVENPAQIREMFD DVSYDKGACILNMLREYLSADAFKSGIVQYLQKHSYKNTKNEDLWDSMASICPTDGVKGMDG FCSRSQHSSSSSHWHQEGVDVKTMMNTWTLQ**K**GFPLITITVRGRNVHMKQEHYMKGSDGAPD TGYLWHVPLTFITSKS**D**MVHRFLLKTKTDVLILPEEVEWIKFNVGMNGYYIVHYEDDGWDSL TGLLKGTHTAVSSNDRASLINNAFQLVSIGKLSIEKALDLSLYLKHETEIMPVFQGLNELIP MYKLMEKRDMNEVETQFKAFLIRLLRDLIDKQTWTDEGSVSE**R**MLRS**Q**LLLLACVHNYQPCV QRAEGYFRKWKESNGNLSLPVDVTLAVFAVGAQSTEGWDFLYSKYQFSLSSTEKSQIEFALC RTQNKEKLQWLLDESFKGDKIKTQEFPQILTLIGRNPVGYPLAWQFLRKNWNKLVQKFELGS SSIAHMVMGTTNQFSTRTRLEEVKGFFSSLKENGSQLRCVQQTIETIEENIGWMDKNFDKIR VWLQSEKLERMGS**<u>ENLYFQS</u>***HHHHHHHHHH* pFASTBAC1-ERAP1_Hap10: <u>MVFLPLKWSLAIMSFLLSSLLALLTVSTPSW</u>CQSTEASPKRSDGTPFPWNKIRLP**E**YVIPVH YDLLIHANLTTLTFWGTTKVEITASQPTSTIILHSHHLQISRATLRKGAGERLSEEPLQVLE HP**P**QEQIALLAPEPLLVGLPYTVVIHYAGNLSETFHGFYKSTYRTKEGELRILASTQFEPTA ARMAFPCFDEPAFKASFSIKIRREPRHLAISNMPLVKSVTVAEGLIEDHFDVTVKMSTYLVA FIISDFESVSKITKSGVKVSVYAVPDK**I**NQADYALDAAVTLLEFYEDYFSIPYPLPKQDLAA IPDFQSGAMENWGLTTYRESALLFDAEKSSASSKL**G**IT**V**TVAHELAHQWFGNLVTMEWWNDL WLNEGFAKFMEFVSVSVTHPELKVGDYFFGKCFDAMEVDALNSSHPVSTPVENPAQIREMFD DVSYDKGACILNMLREYLSADAFKSGIVQYLQKHSYKNTKNEDLWDSMASICPTDGVKGMDG FCSRSQHSSSSSHWHQEGVDVKTMMNTWTLQ**R**GFPLITITVRGRNVHMKQEHYMKGSDGAPD TGYLWHVPLTFITSKS**N**MVHRFLLKTKTDVLILPEEVEWIKFNVGMNGYYIVHYEDDGWDSL TGLLKGTHTAVSSNDRASLINNAFQLVSIGKLSIEKALDLSLYLKHETEIMPVFQGLNELIP MYKLMEKRDMNEVETQFKAFLIRLLRDLIDKQTWTDEGSVSE**Q**MLRS**E**LLLLACVHNYQPCV QRAEGYFRKWKESNGNLSLPVDVTLAVFAVGAQSTEGWDFLYSKYQFSLSSTEKSQIEFALC RTQNKEKLQWLLDESFKGDKIKTQEFPQILTLIGRNPVGYPLAWQFLRKNWNKLVQKFELGS SSIAHMVMGTTNQFSTRTRLEEVKGFFSSLKENGSQLRCVQQTIETIEENIGWMDKNFDKIR VWLQSEKLERMGS**<u>ENLYFQS</u>***HHHHHHHHHH*

### Peptides

Peptides were custom synthesized by JPT Peptide Technologies GmbH (Berlin, Germany), validated by mass spectrometry and were >95% pure as judged by reverse-phase HPLC (chromolithC-18 column, Merck).

### Site-directed mutagenesis

Mutagenesis reactions to introduce the two single mutations into ERAP1 HAP10 WT (V349M and Q725R), as well as the double one (V349M & Q725R), were performed using the Quickchange II kit (Agilent Technologies), according to the manufacturer’s instructions. Primers were designed with the Quick-change Primer Design tool (http://www.genomics.agilent.com). The initial template was the pFastBacI-ERAP1 HAP10 WT plasmid. The sequences of primers were (5ʹ−3ʹ) as follows: ***V349M FW:*** *CAAGCTGGGTATCACCATGACCCTCGCTCACGAAC **V349M REV:** CTTCGTGAGCGACGGTCATGGTGATACCCAGCTTG **Q725R FW:** CGAAGGCAGCGTGTCTGAAAGAATGCTCCGTTCTGAACTC **Q725R REV**: GAGTTCAGAACGGAGCATTCTTTCAGACACGCTGCCTTCG* After each mutagenesis reaction, the selected mutation was verified by sequencing (VBC Genomics, Austria). For generating the double mutant, the Q725R mutant was used as a template to introduce the V349M mutation.

### Construction of recombinant baculoviruses

All recombinant baculoviruses were constructed using the Bac-to-Bac System by Invitrogen Life technologies, according to the manufacturer’s instructions. The plasmids carrying the desired mutations (donor plasmids) were transformed into competent *E. coli* DH10Bac cells which carry the bacmid that encodes for the mature baculovirus. Upon transposition of the gene of interest from the donor plasmid into the bacmid, the recombinant bacmid harbouring the gene of interest is generated. The transformed cells were plated in Luria Broth (LB) agar petri dishes containing the appropriate antibiotics (50 μg/mL kanamycin, 7 μg/mL gentamicin, and 10 μg/mL tetracycline), as well as 100 μg/ml Bluo-gal, and 40 μg/ml IPTG for the blue-white screening. White colonies indicate the clones harbouring the recombinant bacmid with donor insertion. The white phenotype was verified by re-streaking of a colony to a fresh petri dish containing the same selection reagents. One colony was selected and grown overnight in liquid LB cultures with antibiotics to amplify the bacmid. The following day, the culture was collected and centrifuged at 4000 rpm for 20 min. The cell pellet was resuspended in 100 mM Tris (pH 8.0), 8% sucrose, and 10 mM EDTA, in the presence of RNase A (100 μg/mL) and lysozyme (50 μg/mL), and the mixture was incubated at room temperature for 10 min. Cells were then lysed with 1% SDS in Tris-EDTA buffer, and after a 10 min incubation at 37 °C, potassium acetate (5 M) was added to the mixture. After centrifugation to remove the debris, the DNA in the supernatant was precipitated with isopropanol. The pellet was washed with 70% EtOH, air-dried, and resuspended in Tris-EDTA buffer (pH 8.0). The gene’s transposition was verified by PCR, according to the manufacturer’s instructions. The bacmid DNA was then transfected to insect Sf9 adherent cells, grown in SF900II serum-free medium (Life Technologies). For transfection, the Cellfectin reagent (Invitrogen) was used according to the manufacturer’s instructions. The cells were allowed to grow in the 6 well-plates during the following days and once the signs of viral infection were obvious (4-10 days) the supernatant was collected by centrifugation (4500 rpm, 5 min, RT) and filtered using 0.2 μm filters (Merck Millipore). The collected supernatant, comprising the P0 viral stock, was then used to amplify the viral titer by infecting Sf9 insect cells in T75 flasks. Once the cells exhibited significant signs of infection, the culture’s supernatant was collected, comprising the P1 viral stock. This stock was used to infect Hi5 insect cells in suspension cultures to produce the P2 the viral stock, characterized by higher viral titer, which was then used for all protein expression.

### Protein expression and purification

The recombinant enzymes were produced by infecting Hi5 insect cell cultures (50-100 ml) with the P2 baculovirus stock, in a 1:25-1:50 v/v ratio. The culture’s supernatant was collected 72h post-infection. All pFastBacI constructs used to express ERAP1 allotypes contain a C-terminal 10His-tag, which allows protein purification by Immobilized Metal Affinity Chromatography using Ni-NTA Sepharose. Cell supernatant is incubated with the Ni-beads in the presence of 20mM imidazole in phosphate buffer at pH=7.0, washed and protein eluted using increasing concentrations of imidazole buffers. Fractions of 1-1.5 ml volume were collected, and the enzymatic activity was measured using a kinetic assay that follows the hydrolysis of a small fluorogenic substrate (see below). The fractions containing the purified enzyme were dialysed overnight against a buffer containing 10 mM HEPES, 100 mM NaCl pH=7.0 or buffer exchanged using a HiTrap 5ml desalting column (Cytiva). The final concentration of the purified recombinant enzyme was calculated by densitometry on SDS-PAGE, using a protein of known concentration as a reference.

### Fluorogenic enzymatic assay

The aminopeptidase activity of the recombinant ERAP1 was measured by following the change in the fluorescence signal produced upon hydrolysis of the fluorescent substrate L-Leucine-7-amido-4-methylcoumarin (L-AMC, Sigma-Aldrich) or the change in absorbance during the hydrolysis of the chromogenic substrate L-leucine-p-nitroanilide (L-pNA, Sigma-Aldrich). In the first case, the fluorescence was measured at 460 nm, whereas the excitation was set at 380 nm. In the second case, the absorbance was measured at 405 nm. Measurements were performed on a TECAN infinite M200 microplate fluorescence reader or the microplate reader Synergy H1 by BioTek. The buffer used consisted of HEPES 20mM pH=7.0, NaCl 150mM and Tween20 0.002%. The specific enzymatic activity (mol product (AMC)/mol enzyme/sec) was calculated using a standard curve for AMC. The enzymatic activity was calculated by the slope of the kinetic gradient.

To measure the activation of small substrate trimming by GSK849, the change in the fluorescent signal produced upon digestion of the substrate L-AMC was also followed. A threefold dilution series (1/3log scale) of the modulator was made in reaction buffer consisting of 20mM HEPES pH=7, 150mM NaCl and 0.002% Tween20, containing a final concentration of 1% DMSO, upon dilutions of the stock solution of GSK849 (in 100% DMSO) in the reaction buffer. The dilution series of the compound was put in duplicates in a 96-well black plate. The enzyme/substrate mix was prepared in the reaction’s buffer so that the substrate’s final concentration in the reaction would be 50 μM and the enzyme concentration 10 nM for HAP1 and HAP10 variants and 50 nM for wild-type HAP10. The enzymatic activity was calculated for every concentration point from the slope of the kinetic data, normalized to control (1% DMSO) and plotted in Graph Pad Prism 8.0, by applying the log(agonist) vs. response -- Variable slope (four parameters) non-linear fit.

### Peptide trimming followed by HPLC

10 μΜ of peptide was used for each digestion reaction, in a final reaction volume of 100 μl in a buffer containing 50mM HEPES, pH=7, and 150mM NaCl. For the digestion of the 9-mer peptide, the enzymes were tested at the following concentrations: 4 nM for HAP1, 10 nM for HAP10 variants and 20 nM for HAP10 wild-type. For the digestion of the 10-mer peptide, the enzyme concentration used was 1 nM for HAP1, 2nM for HAP10 variants and 10 nM for HAP10. All reactions were performed in triplicates and incubated for 1 hr at 37 °C in a water bath, upon the addition of the enzyme. After the incubation, the reactions were quenched by adding 100 μl of 1% trifluoracetic acid (TFA) solution and kept on ice until analysis. The reactions were analysed by reversed phase HPLC (Chromolith C-18 column, Merck) by following the absorbance at 220 nm. A linear gradient elution system was used (solvent A, 0.05% TFA and 5% ACN; solvent B, 0.05% TFA and 50% ACN). The reaction’s progression was calculated by integrating the area under each peak. The percentage of the area of the product’s peak (P) to the sum of the area of the remaining substrate (S) plus the product (P) peaks, (P/P+S*100%) was used to calculate the mol of product produced which was used to determine the specific enzymatic activity.

### Michaelis-Menten analysis

Michaelis Menten analysis was carried out for the hydrolysis of the small chromogenic substrate L-pNA, by following the reaction’s rate for different substrate concentrations (0.1 – 6 mM). Serial dilutions were prepared in Eppendorf tubes at an intermediate concentration (3 times higher than the final desired concentration) and 50 μl of each concentration point were put in a 96-well plate in duplicates. The enzyme was diluted in the reaction buffer (20mM HEPES pH=7.0, 150mM NaCl, 0.002% Tween20), so that the final concentration upon addition of 100 μl in the well, would be 10 nM. All measurements of absorbance were taken in the microplate reader Synergy H1 of Biotek at 405 nm. The specific enzymatic activity was calculated for every substrate concentration by calculating the slope of the kinetic data. The specific activities were fit to a classical Michaelis Menten or an allosteric (cooperative) model, depending on the quality of fit, using GraphPad Prism software. For the allosteric model, the fit was performed using the equation Y = (V_max_ * X*h)/(K_prime_ + X*h), where X is the substrate concentration, Y the initial reaction rate, V_max_ is the reaction rate at infinite time, and h the Hill slope.

To define the mechanism of action of the phosphinic pseudo-peptide transition state analogue DG013A, the Michaelis Menten experiment was performed as described above, with the addition of different concentrations of the inhibitor: 10, 20 and 30 nM (final concentrations). The inhibitor was added to the enzyme/buffer solution before the addition of 100ul in the plate, already containing 50 μl of the substrate’s serial dilutions.

### Eyring Analysis

The enzymatic rate for the hydrolysis of L-AMC by ERAP1 variants was measured at 6 different temperature points: 25, 28, 31, 34, 37 and 40 °C for HAP1 and HAP10’s variants and 28, 31, 34, 37, 40 and 43 °C for HAP10 wt. All measurements were carried out in triplicates at the microplate reader Synergy H1 of BioTek. For HAP1, and HAP10 variants, 50 μΜ of L-AMC and 10 nM of enzyme were used as final concentrations in a 150 μl final volume. For wild-type HAP10 a final concentration of 50 nM of enzyme was used. Before each measurement, the substrate, enzyme solutions and the measurement plate were preincubated at the required temperature for 10 minutes. The specific enzymatic rate was calculated for each temperature as described before. For Eyring analysis, the data were fit in Graph Pad Prism according to the equation:

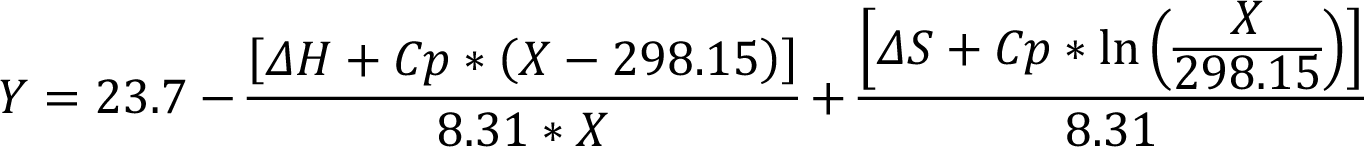

where: Y= ln(k_cat_/T) and X=temperature expressed in Kelvin (K), ΔH and ΔS correspond to the enthalpy and entropy of activation respectively, and C_p_ is the heat capacity of activation. The value 23.7 corresponds to ln(kb/h), where kb is Boltzmann’s constant and h is Plank’s constant. The value 8.31 is the gas constant, R and 298.15 corresponds to the reference temperature, T_0_=25°C (298.15K).

### Effects of solvent viscosity to enzymatic turnover rate

To characterize the viscosity effect of the solvent on the enzymatic activity of ERAP1 variants, the turnover rate (*k*_cat_) of the enzymatic reactions for the cleavage of L-pNA was measured in buffers with various glycerol concentrations, at 25°C. Four different glycerol concentrations were tested: 0, 9, 18 and 27% (v/v). Michaelis Menten analysis was carried out to determine the turnover rate in each case, as described in the corresponding Methods section. The concentration of enzymes used was 10 nM for Hap1 and for Hap10 carrying the double mutation (V349M & Q725R), 30 nM for wild-type Hap10 and 15 nM for the single mutation carrying variants (V349M, Q725R). The ratios kcat(0)/kcat(η), where kcat (0) and kcat(η) are the turnover rates in the absence and presence of glycerol respectively, were plotted as a function of the relative viscosity(η_rel_) at the tested temperature, calculated based on previously reported viscosity values of aqueous solutions^20^. The data were fitted in Graph Pad Prism using equation (2): kcat(0)/kcat(η)= m*(η_rel_-1)+1, where m is the slope and represents the degree of the rate dependency on viscosity.

## RESULTS

To investigate the relative contribution of the polymorphic positions 349 and 725 to the unusual functions of ERAP1 allotype 10, we used site-directed mutagenesis to generate ERAP1 allotype 10 variants V349M and Q725R as well as the double variant V349M/Q725R, expressed the recombinant proteins in insect cells after infection with the appropriate recombinant baculovirus and purified them by affinity chromatography as described in the methods section. Figure 1A, shows a schematic representation of the locations of the mutated residues relative to the ERAP1 active site and Figure 1B, SDS-PAGE analysis of the ERAP1 variants generated for this study. The sequences of the constructs used are shown in the Methods section.

**Figure 1:**
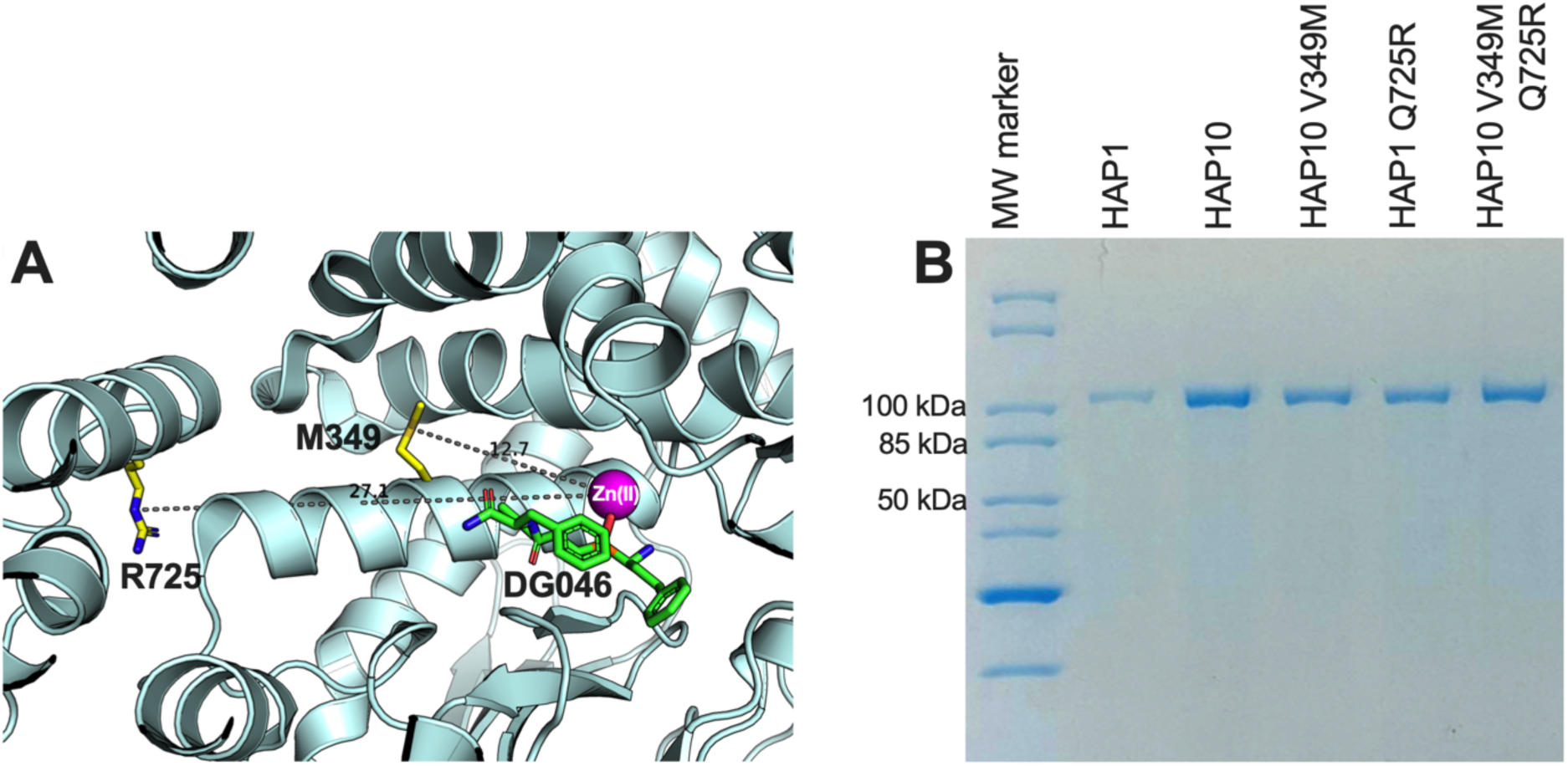
**Panel A**, schematic representation of the ERAP1 crystal structure of allotype 1 (PDB ID 6Q4R^21^) showing the active site where the catalytic zinc(II) atom is located (shown as a magenta sphere) and bound pseudopeptidic inhibitor (depicted as green sticks) as well as the location of the two polymorphic locations that are unique to ERAP1 allotype 10 (shown in yellow sticks). The dotted lines indicate the distance (in Å) of the polymorphic amino acids 349M and 725R from the active site (12.7Å and 27.1Å away from the catalytic zinc atom, respectively). **Panel B**, SDS-PAGE showing the purified ERAP1 allotypes and variants used in this study.

To evaluate the role of ERAP1 allotypes and specifically the role of polymorphic positions 349 and 725 on catalytic rates, we used the recombinant ERAP1 variants shown in Figure 1B to follow the trimming of three model peptides. We chose peptides of different lengths since ERAP1 peptide trimming is length dependent^22^. Thus we tested: i) the 9mer antigenic peptide YTAFTIPSI from the Gag-Pol polyprotein of the Human Immunodeficiency Virus 1 ^23^, ii) the 10mer precursor LRVYEKMALY of an HLA-A*03 ligand and iii) the 11mer precursor VSVRSRRCLRL of the antigenic peptide VRSRRCLRL from the human ADAMTS-like protein 5^24^ (Figure 2). The specific activity for the trimming of the 9mer peptide varied by almost 100-fold between the two ERAP1 allotypes, consistent with previous observations^14^, and the widely held notion that allotype 10 is a sub-active, or non-functional allotype. Mutating positions 349 and 725 of allotype 10 to the corresponding amino acids found in allotype 1 resulted in ERAP1 variants with enhanced activity, about 10-fold for each mutation and up to 20-fold for the double mutant, although the variants were still significantly less active than allotype 1. Strikingly, when testing the trimming of a 10mer substrate, the situation was reversed and allotype 10 was no longer less active but rather presented up to 2-fold higher activity. The mutations resulted in a decrease of activity and the double mutant was found to be about as active as allotype 1. When trimming an 11mer peptide, allotype 10 was about 2-fold less active than allotype 1 and the mutations resulted in an additive rescuing of this loss in activity. These experiments suggest a complex framework of activities for each ERAP1 allotype that is consistent with results previously obtained from single-point kinetic analysis of a peptide library derived from the SARS-CoV-2 S1 spike glycoprotein^19^. Our results do, however, highlight that the allotype 10 specific SNPs at positions 349 and 725 are important for shaping the activity of ERAP1 in particular for the trimming of 9mer antigenic peptides. This complex landscape of substrate-dependent activity is a hallmark of ERAP1^25^ and is likely due to extensive atomic interactions between the peptide side-chains and the substrate cavity of the enzyme as demonstrated by x-ray crystallography^26^, which are highly dependent on the length and the sequence of the peptide.

**Figure 2:**
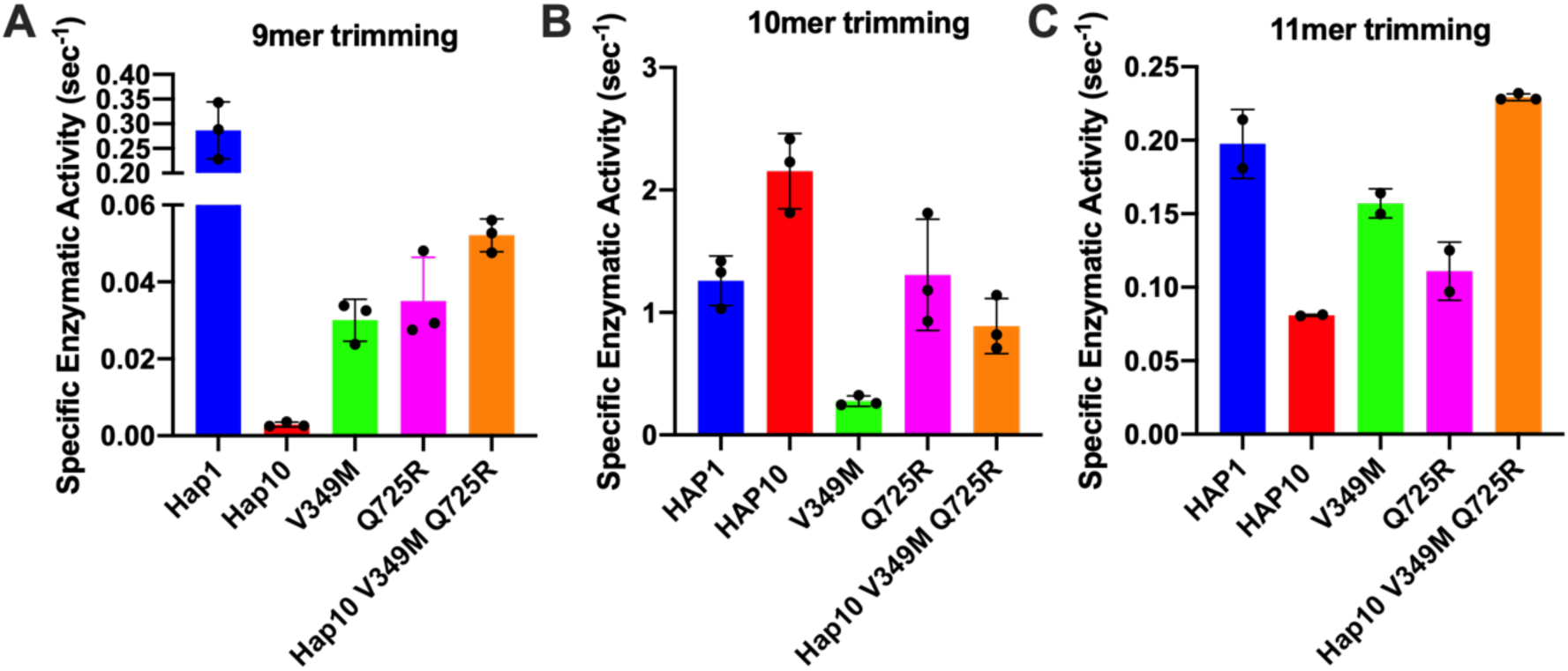
Specific enzymatic activity for the trimming of the N-terminal residue of three model peptides by ERAP1 allotypes 1 and 10 as well as variants of ERAP1 allotype 10. **Panel A,** peptide YTAFTIPSI; **Panel B,** peptide LRVYEKMALY; **Panel C,** peptide VSVRSRRCLRL.

To further examine the functional differences between ERAP1 allotypes and polymorphisms at locations 349 and 725 we performed a series of kinetic and thermodynamic analyses using typical dipeptide substrates. We chose to focus our analysis on dipeptide substrates to minimize the well-established effect of peptide sequence on ERAP1 enzymatic kinetics which complicates interpretation and instead put emphasis on the events occurring in the catalytic centre of ERAP1. We first measured specific rates of hydrolysis for substrates L-AMC (fluorogenic) and L-pNA (chromogenic) (Figure 3A and 3B). In both cases allotype 1 was more active compared to allotype 10, and more specifically 38-fold more active for L-AMC and 8-fold more active for L-pNA. Introduction of the mutations V349M and Q725R, partially rescued the reduced activity of allotype 10, although this was more clear in the L-pNA assay in which the effect of the mutations was additive. Overall, the relative activities between allotypes and mutants resemble the motif seen for the 9mer peptide but not for larger peptides, consistent with previous observations that 9 amino acids is an important length cutoff for the activity of ERAP1^22^.

**Figure 3:**
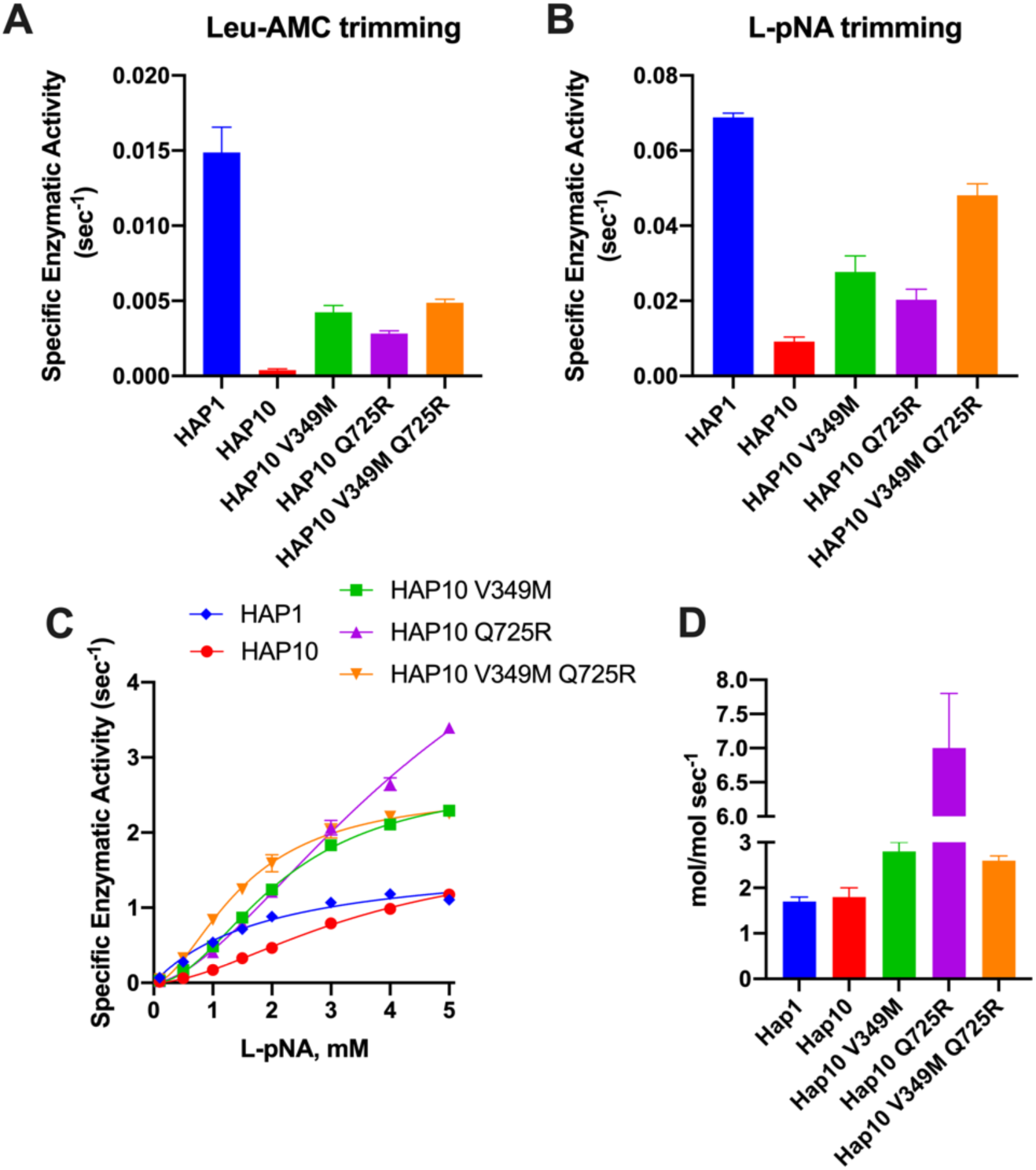
Specific activity of trimming of the model dipeptidic substrates Leu-AMC (**Panel A**) and Leu-pNA (**Panel B**) by ERAP1 allotypes and variants at positions 349 and 725. **Panel C,** Michaelis Menten analysis of ERAP1 allotypes and variants using the model dipeptidic substrate Leu-pNA.

Michaelis-Menten (MM) analysis further highlighted the differences between the two allotypes (Figure 3C). The dependence of catalytic activity on substrate concentration for allotype 1 fits a classical MM model with a calculated *k*_cat_ of 1.7±0.1 sec^-1^. In contrast, allotype 10 can only be fit to an allosteric sigmoidal model with a calculated *k*_cat_ of 1.8±0.2 sec^-1^. This indicates that the observed differences between these two allotypes are ameliorated at high substrate concentrations and that the low apparent catalytic turnover of allotype 10 is not due to overall lower catalytic efficiency but rather due to a requirement for allosteric activation by the substrate. Interestingly, introduction of the V349M and Q725R mutations to allotype 10, did not result in a change of the allosteric behaviour, but did result in a much higher *k*_cat_, especially for the Q725R variant (Figure 3D). Overall, MM analysis suggested that the relative contribution of allotypic variation to enzyme activity is highly dependent on substrate concentration, an important parameter that has not been evaluated before. Given that the substrate concentrations of ERAP1 in the cell are not known, this finding suggests that the interpretation of allotypic differences *in cellulo* may be heavily influenced by the unknown local concentrations of the peptides being analyzed.

Given the significant mechanistic differences revealed by MM analysis, we used a transition state analogue to further probe the catalytic mechanism of ERAP1. DG013A is a pseudophosphinic tripeptide that is a transition-state analogue for metallo-aminopeptidases, optimized for ERAP1 and homologous ERAP2 and IRAP ^27^, which has been extensively used as a tool compound for inhibiting ERAP1 in several systems^28^. A crystal structure of DG013A with ERAP1 is available and revealed a canonical binding mode resembling the tetrahedral intermediate of the LTA4 hydrolase catalytic mechanism ^29,30^. This binding mode would be consistent with competitive inhibition kinetics. Indeed, MM analysis of the effect of DG013A on the kinetic parameters of allotype 1 suggested a major effect on the K_M_, consistent with competitive inhibition (Figure 4A, 4F). Surprisingly, the same analysis for allotype 10 revealed a non-competitive inhibition mechanism with major effects on the *k*_cat_ of the enzyme (Figure 4B, 4F). Although at first glance this may appear to contradict with the binding site of the inhibitor, this unusual behaviour has been reported before for systems where multiple conformations are in equilibrium with each other^31,32^ something previously proposed for ERAP1^33^. Furthermore, the allosteric behaviour of the MM curve was not affected by the presence of the inhibitor. The introduction of the V349M or Q725R mutations did not affect this allosteric behaviour, nor the non-competitive nature of the inhibition. This analysis thus concludes that allotypes 1 and 10 present significant mechanistic differences in how they recognize the transition state of small substrates and that these differences are not determined by the polymorphisms at positions 349 and 725.

**Figure 4:**
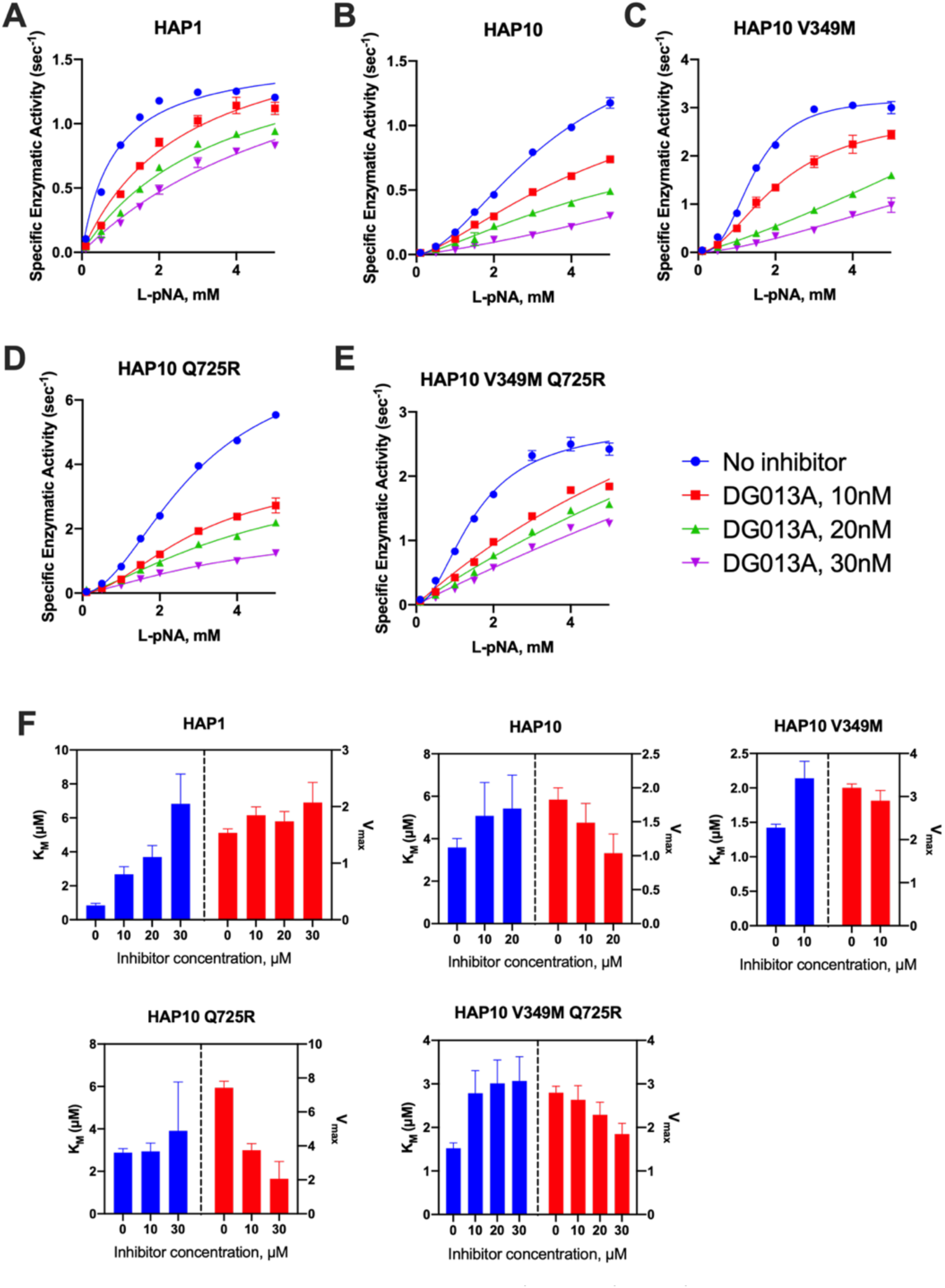
**Panels A-E**, Michaelis-Menten analysis of the effect of the transition-state analogue inhibitor DG013A on the kinetics of trimming of Leu-pNA by ERAP1 allotypes and variants. **Panel F,** calculated enzymatic parameters K_M_ and V_max_ from the fits in panels A-E.

Motivated by the surprising behaviour of the transition-state analogue, we decided to probe the regulatory site of ERAP1 by utilizing a previously characterized allosteric modulator of ERAP1, named GSK849^34^. This compound was discovered by a high-throughput screening campaign as an ERAP1 activator, although it is a competitive inhibitor for longer peptides^34^. GSK849 binds to the regulatory site of ERAP1, which is located more than 30Å away from the catalytic center, a site that can accommodate the C-terminus of many peptide substrates and can allosterically enhance ERAP1 activity through networks of correlated residue motions^26^. Titration of GSK849 while following L-AMC hydrolysis resulted in about 3.5-fold activation of allotype 1 (Figure 5A). Activation of allotype 10 was only evident at higher concentrations of GSK849 and reached similar levels as allotype 1, although it was not possible to clearly define a plateau at achievable compound concentrations. Surprisingly, both mutations resulted in much higher levels of activation of up to 15-fold. We conclude that the polymorphic positions 349 and 725 are important for the communication between the regulatory and active sites.

**Figure 5:**
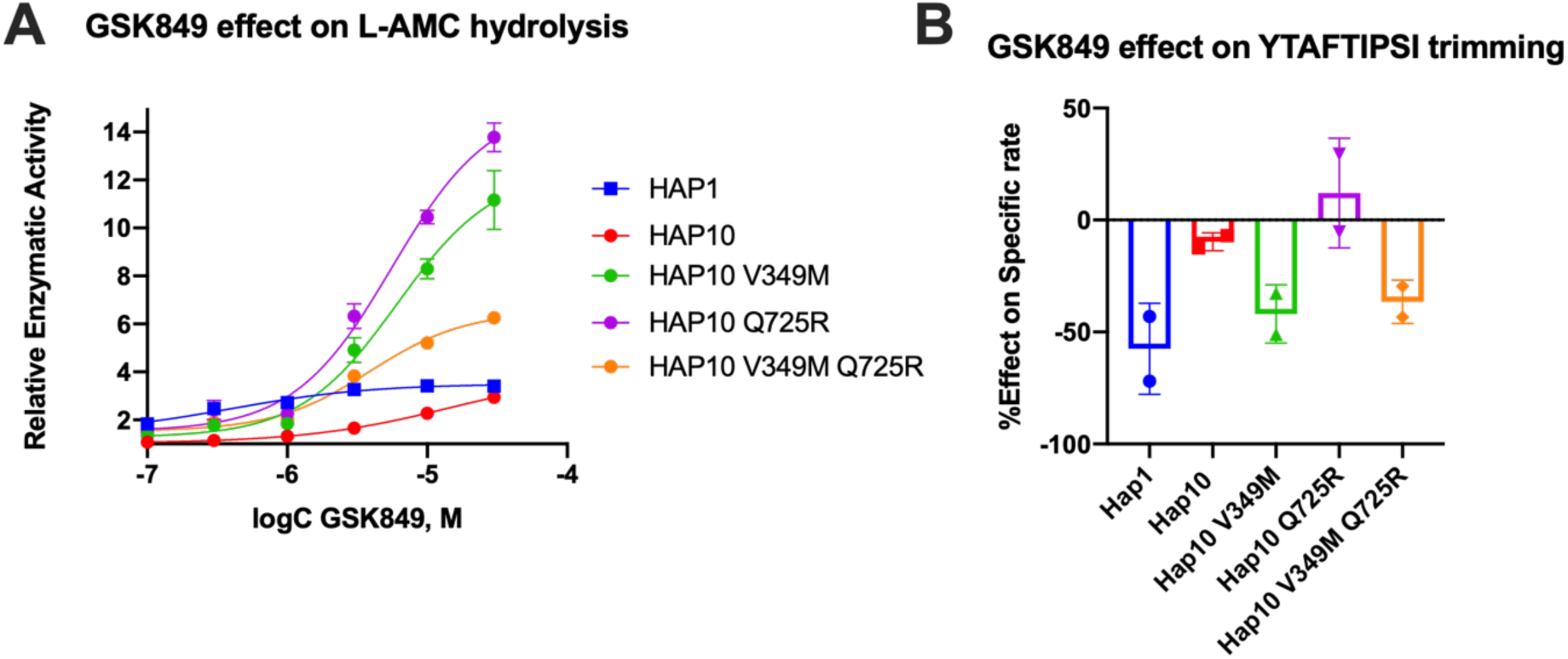
**Panel A**, effect of the allosteric inhibitor GSK849 on the activity of ERAP1 allotypes and variants when following the trimming of the dipeptidic substrate Leu-AMC. **Panel B**, effect of GSK489 on the specific rate of ERAP1 allotypes and variants for the trimming of the 9mer antigenic epitope YTAFTIPSI.

Given the large effect of these polymorphic positions on the activation of ERAP1 by this allosteric modulator, we also examined its effect on the trimming of the 9mer peptide shown in Figure 2A. In Figure 5B, we show the % inhibitory effect of 10 μΜ GSK849 on the trimming of the N-terminus of the peptide YTAFTIPSI by ERAP1 allotypes and mutants. The inhibitor was effective in blocking most of the trimming by allotype 1, but had little effect on allotype 10, as previously shown^14^. The V349M mutation recovered the ability of GSK849 to inhibit ERAP1 allotype 10, whereas Q725R had no effect; the double mutant mirrored the effect of the V349M polymorphism. Overall, we find that the V349M polymorphism but not the Q725R affected the inhibition capability of GSK349, suggesting that the two polymorphisms have different effects on the communication between the regulatory and active sites when using longer peptidic substrates.

Given the surprising extent of differences in kinetic parameters between ERAP1 allotypes we hypothesized that these changes may be due to changes in the thermodynamics of the catalytic reaction. To test this hypothesis, we measured the temperature-rate dependency of the enzymatic reactions by performing Eyring analysis for the hydrolysis of Leu-AMC. This analysis allows the determination of the thermodynamic properties of the system, namely the enthalpy of activation *ΔΗ*^‡^ and the entropy of activation *ΔS*^‡^. Eyring plots follow the ln(k_cat_/T) as a function of temperature, which usually corresponds to a straight line^35^. The slope of this line is proportional to the enthalpy of activation *ΔΗ*^‡^, and the intercept is related to the entropy of activation *ΔS^‡^*. Indeed, this was observed for allotype 1, as shown in Figure 6A. The linear relationship implies that the enthalpy of activation and the entropy of activation are constants over the temperature range studied. In contrast, we observed significant deviations in linearity for allotype 10 and its V349M or Q725R variants (Figure 6A) which indicates that *ΔΗ*^‡^ and *ΔS*^‡^ may vary with temperature. This behaviour has been interpreted to indicate a non-zero heat capacity of activation (*ΔCp*^‡^), a statistical thermodynamic property that describes the temperature dependency of *ΔΗ*^‡^ and *ΔS*^‡^, and expresses the difference in the system’s heat capacity between the transition and the ground states^36^. All these three parameters (enthalpy, entropy and heat capacity of activation) were determined by fitting the data to eq (1), as described in the Methods section and the results are shown in Figure 6B-D.

**Figure 6:**
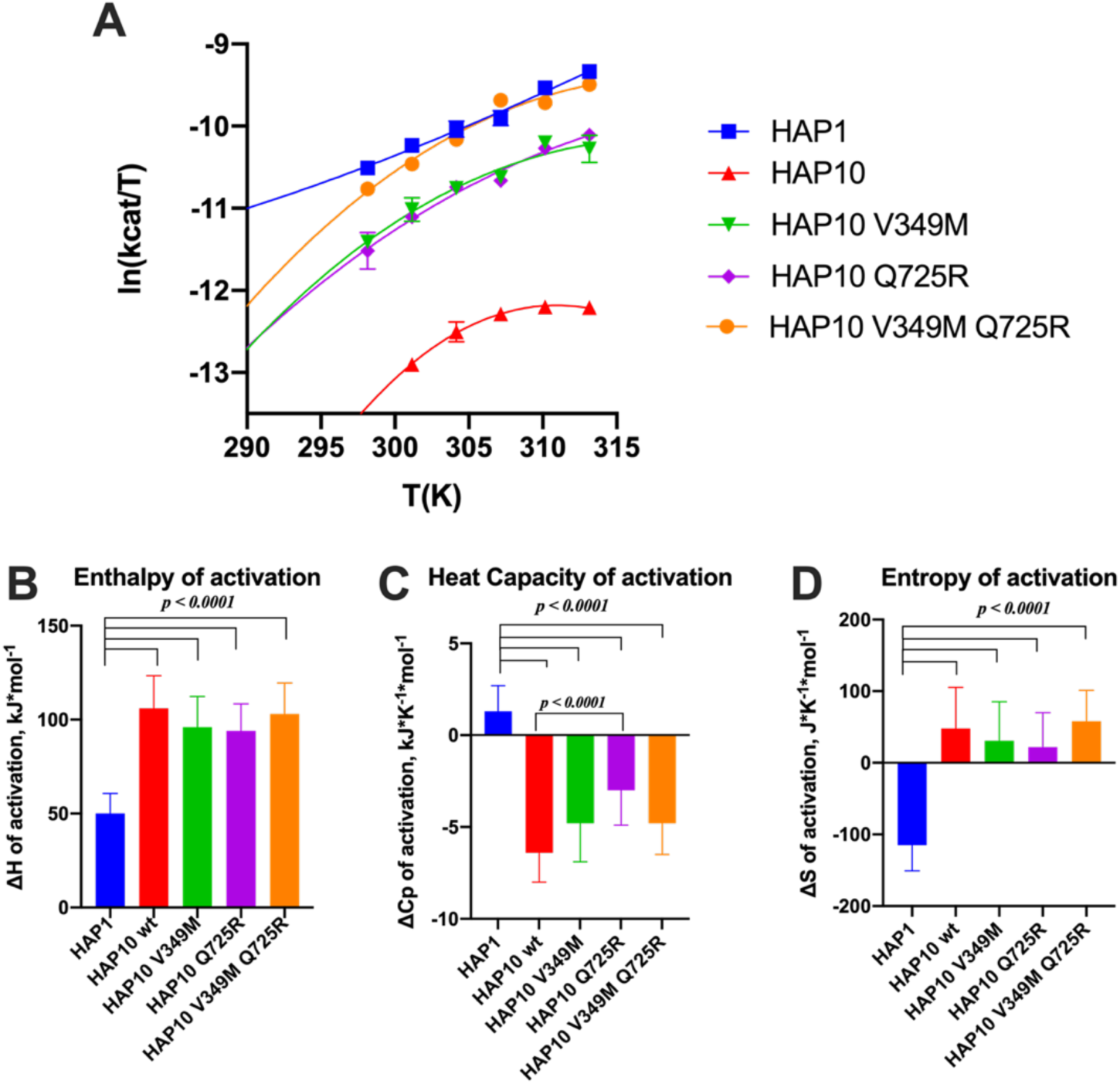
**Panel A**, effect of temperature on the activity of ERAP1 allotypes and variants. Curves were fit to the Eyring equation as described in the Methods section. **Panels B-D,** calculated parameters for the enthalpy of activation, heat capacity of activation and entropy of activation based on the fits shown in panel A.

Overall, this analysis indicated major thermodynamic differences between allotypes 1 and 10 with only a limited contribution of positions 349 and 725. In particular, allotype 10 and its variants exhibited a much higher enthalpy and entropy of activation and a highly negative heat capacity of activation. Negative *ΔCp*^‡^ values have been reported for enzymes when the chemical reaction is rate-limiting, resulting in non-linear Eyring plots^37^. The surprisingly large differences between allotype 1 and allotype 10 and variants suggest a complex reaction pathway with multiple stages in which the observed rate constant is dominated by the slowest stage, which can vary between allotypes.

Diffusion can be a rate-limiting step for some enzymatic reactions. To explore whether diffusional steps influence the enzymatic reaction catalyzed by ERAP1 and if these are affected by allotypic variation, we determined the solvent viscosity effect on the enzymatic rate by performing a Kinetic Solvent Viscosity Effect analysis (KSVE). KSVE analysis gives insight on whether the reaction is bottlenecked by diffusion-controlled steps, starting at initial substrate association throughout product release^38^. We calculated the ratio of *k*_cat_ compared to a reference viscosity and plotted the results as a function of viscosity (Figure 7A). For all ERAP1 allotypes tested, there was a large positive slope suggesting that the ERAP1-catalyzed reaction is strongly influenced by diffusion (Figure 7B). The calculated slope was higher for allotype 10 compared to allotype 1, but it was greatly reduced in an additive fashion by the introduction of the mutations V349M and Q725R. Thus, we conclude that ERAP1 catalytic turnover is bottlenecked by a step that is highly dependent on solvent viscosity and this is greatly affected by the allotype 10 polymorphisms V349M and Q725R. A positive slope in the KSVE graph has been observed in cases where the bottleneck is product release. Given that product release from the closed ERAP1 conformation requires a substantial conformational change (the closed conformation does not provide direct access to the external solvent) we interpret this result to suggest that this allotypic variation affects the conformational opening of ERAP1.

**Figure 7:**
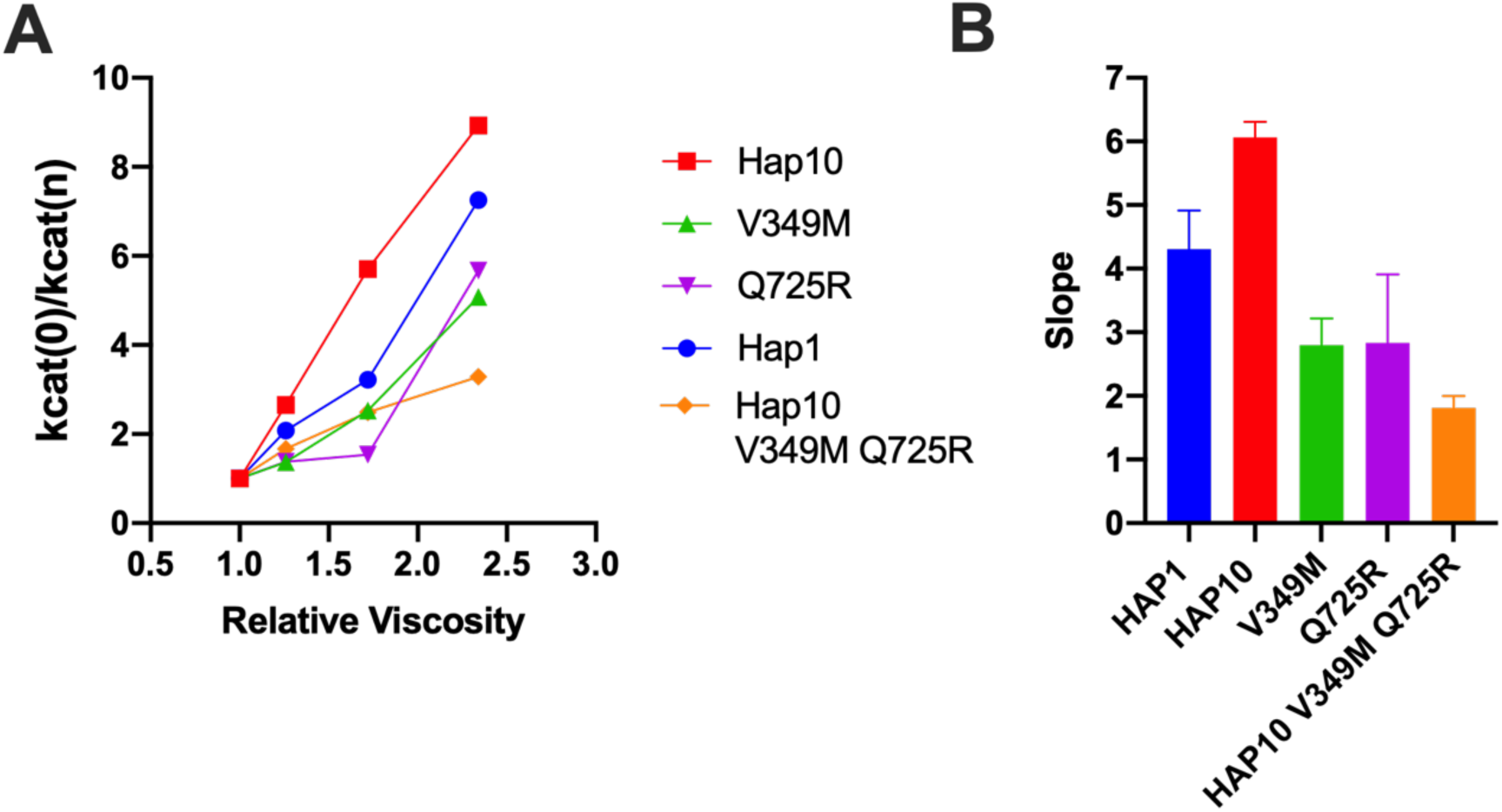
**Panel A**, Solvent viscosity effect on the enzymatic rate of ERAP1 variants. The Kinetic Solvent Viscosity Effect (KSVE) curves depicted are the result of the fit described in the methods section. It follows the *k*_cat_ ratios in function of different glycerol concentrations in the reaction’s buffer. **Panel B,** slope values of the KSVE curves fit for ERAP1 variants, representative of the solvent viscosity effect.

## DISCUSSION

### Polymorphic variation in ERAP1

Several SNPs in ERAP1 have been associated with predisposition to disease, most notably autoimmunity but more recently also with cancer, and in many cases, this association comes in epistasis with specific HLA alleles that were already known to predispose to disease, suggesting a functional interaction^10,39^. Indeed, functional effects of several SNPs are related to antigenic peptide trimming, thus providing a mechanistic link to disease pathogenesis through aberrant antigen presentation, albeit other mechanisms, including ER stress and protein misfolding, have also been proposed^40–43^. A hallmark of ERAP1 SNPs is that they are all located away from the active site and are found in particular sets (allotypes) within the population^13,14^. Recent analyses suggest that these allotypes consist of SNPs that synergize to give rise to functional properties. The most notable is allotype 10, which is shared amongst 22.4% of Europeans, associates with a protective phenotype for autoimmunity and has been shown to be sub-active against some peptide substrates^15^. More detailed analysis however revealed that allotype 10 carries distinct substrate specificity^14^ which is very difficult to decipher due to the complicated enzyme-substrate interactions for ERAP1^26^. Thus, unravelling the mechanisms behind the functional properties of allotype 10 is crucial for a deep understanding of the role of ERAP1 in the development of autoimmunity as well as in adaptive immune response to cancer.

### ERAP1 conformational transitions are part of the catalytic cycle

ERAP1 has been crystallized in two distinct conformations, termed the “open” and the “closed” which differ on the accessibility of the catalytic site and of the large internal cavity to the external solvent. The closed conformation has been proposed to feature a more structurally optimized catalytic site, resulting in higher catalytic rates and the open conformation has been proposed to be important for initial substrate capture, especially for large peptide substrates. The transition between these conformations is likely to be central to the ERAP1 catalytic cycle^29^ and molecular dynamics calculations have suggested a relatively shallow energetic landscape that can be modified by common SNPs^33,44^. A possible explanation for the experimental observations described here is that combinations of SNPs as found in allotypes, such as allotype 10, synergize to shape this energetic landscape, affecting the conformational equilibrium and dynamics to regulate turnover rates. The interplay of this energetic landscape with substrates can also further finetune substrate-specific activities. Given the results presented here, a more thorough re-examination of ERAP1 molecular dynamics in the context of specific allotypes and substrates is warranted to provide a more solid structural framework for understanding their functional repercussions.

### Fundamental mechanistic differences between ERAP1 allotypes suggest alterations in rate-limiting steps

Our results suggest highly significant mechanistic differences between the ancestral allotype 1 of ERAP1 with the autoimmunity-protective allotype 10. Allotype 10, is much less active but only for shorter substrates (9mers and dipeptides), exhibits allosteric kinetics, reduced communication between the regulatory and active sites, non-competitive kinetics of inhibition by a transition state analogue and very high viscosity dependence. Furthermore, thermodynamic analysis revealed an opposite motif in enthalpy, entropy and heat capacity of activation. Τhe higher absolute value of the activation enthalpy, coupled with the lower absolute value of the activation entropy exhibited by allotype 10, suggests that the catalytic step is characterized by a higher free activation energy, according to the Gibbs equation. This finding aligns with the lower enzymatic rates for the processing of small substrates observed in this allotype. These observations suggest that the catalytic processing of small substrates for allotype 10 is fundamentally different than for allotype 1. This, at first examination, appears paradoxical. The peptide bond cleavage catalysis by Zn(II) aminopeptidases is well established and the mechanism used by LTA4 hydrolase has been proposed to apply also for ERAP1^30^. It would be highly surprising if SNPs located away from the active site directly affect catalytic mechanisms on the atomic level such as substrate hydration effects as suggested by the altered heat capacity of activation^35^. Rather, a multi-step mechanism comprising several conformational changes that generate specific kinetic bottlenecks may constitute a better framework for understanding these effects. In other systems, high dependence of the reaction rates on viscosity has been interpreted to suggest that product release is rate-limiting^38^. Indeed, product release from the closed conformation can be easily envisaged to be a rate-limiting step in ERAP1 since it would require a transition to a more open conformation. Thus, a switch from one rate-limiting step to another between ERAP1 allotypes may result in significant differences in experimentally measured kinetics and thermodynamics since the experiments may be reporting a different step in the catalytic pathway that is rate-limiting. The relatively flat and shallow energetic landscape described for the ERAP1 conformational equilibrium can empower this effect considering that small energy changes can easily reshape this landscape by altering local minima^29,33^.

### Role of polymorphic positions at 349 and 725

The analyses presented here suggest a clear but not absolute role of the allotype-specific polymorphic positions 349 and 725 in shaping allotype 10 enzymatic properties. These two SNPs appear to largely regulate ERAP1 activity and, at least partially, be responsible for the low activity of allotype 10 for smaller substrates. On the contrary, these positions do not appear to define the allosteric behaviour of allotype 10 which may be mediated by other positions that affect mechanism such as K528R^11^. Interestingly, these positions appear to influence the cooperation between ERAP1’s regulatory and active sites and regulate viscosity dependence by affecting product release kinetics. While the atomic contributions of these effects may not be directly obvious, a close inspection of the atomic neighbourhood of these positions in the crystal structures of the open and closed conformation of ERAP1 can provide some insight (Figure 8). Both these polymorphic positions are adjacent to side chains that switch configurations during the transition from open to closed conformation and reversely. Specifically, R725 is within 6Å of K338, a residue that drastically switches position upon transition to the closed conformation of ERAP1 (Figure 8A and B). During this transition, these two charged side chains would come into close proximity, forming repulsive ionic interactions that may increase the energy barrier and alter the equilibrium between the two conformations, promoting the closed conformation. In contrast, the Q725, which is carried exclusively by allotype 10, lacks the positive charge thus negating this interaction. This could result in the enzyme favouring more open states, enabling the more efficient processing of longer peptides. Similarly, M349 makes atomic interactions with Y399 that flips to a different configuration upon the open-to-closed transition (Figure 8C and D). Again, substitution by the much shorter V349 would diminish this interaction altering the conformational equilibrium. While these insights are useful for establishing a framework for the enzymatic effects of these SNPs, a more detailed molecular dynamics study at the atomic level will be necessary to help us gain a deeper understanding of the repercussions of these amino acid substitutions, which is a goal for a future study by our group.

**Figure 8:**
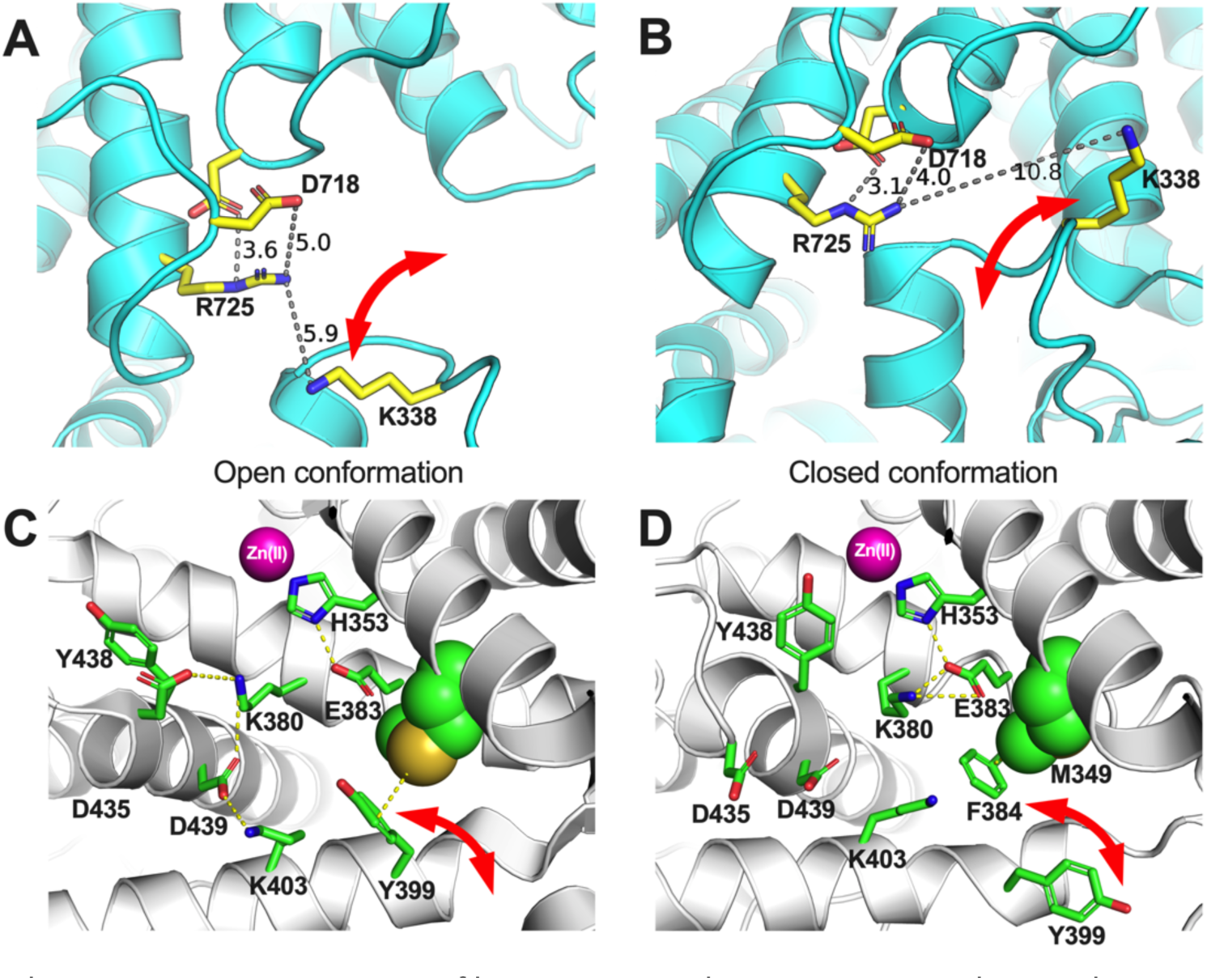
Schematic representations of key amino acids near ERAP1 polymorphic positions 725 (panels A and B) and 349 (panels C and D) in the two known conformations of ERAP1, the open (panels A and C) and the closed (panels B and D). Red arrows indicate amino acid side chains that undergo significant translocations between the two ERAP1 conformations.

The results presented here delineate a framework for understanding the role of ERAP1 allotype in antigenic peptide selection in health and disease. Our data stress that allotype 10, is not a loss-of-function allotype as sometimes described in the literature ^15–18^ but rather an allotype with unique substrate interactions stemming from changes in the conformational dynamics of the enzyme which help define rate-limiting steps of the catalytic cycle. Of particular biological interest is the consistent finding that allotype 10 has a much lower enzymatic activity for smaller substrates. ERAP1 has been described to specialize in trimming extended precursors of antigenic peptides that are longer than 10 amino acids in length ^22^, by employing a molecular ruler mechanism that relies on concurrent occupation of the regulatory or adjacent sites and the active site ^26,29^. While this mechanism helps spare smaller peptides from over-trimming and essential destruction, it is not absolute and ERAP1 can indeed destroy many shorter antigenic peptides. In addition, ERAP1 can contribute to peptide homeostasis in the ER by recycling peptides that do not bind onto MHCI. Thus, an ERAP1 allotype with low activity against shorter peptides while retaining the activity versus larger precursors may provide additional time for some antigenic peptides to bind onto nascent MHC-I, while at the same time leading to an increase of free peptide concentration in the ER. This can result in changes in the repertoire of presented peptides (the immunopeptidome) by favouring the binding of length-optimized peptides, which can reduce antigenicity stemming from longer-length, less stably bound, peptides^45^. At the same time, altered ER peptide homeostasis can affect ER protein folding, especially for the more unstable MHC-I alleles such as the Ankylosing-Spondylitis predisposing HLA-B27, or affect the ER stress by altering unfolded protein responses^40^. In either case, allotype 10 is sufficiently distinct in properties to constitute a potential product of host-pathogen balancing selection in human populations ^6,46^. The interpretation of our findings in the context of the functioning of the human immune system *in vivo* should be performed with care due to specific limitations of our work, as the result of study design. We focused on the *in vitro* characterization of small substrate trimming by the enzymes to understand their underlying basic kinetic and thermodynamic properties. While this was necessary to dissect basic enzyme properties from complex enzyme-substrate interactions important for large substrate selection, these properties may be modified due to atomic interactions with some larger substrates and may have to be re-evaluated in that context. Furthermore, interpolating the effects of allotypic variation on enzymatic function inside the cell assumes a similar expression level for ERAP1 allotypes. This may not be always be the case and indeed previous studies have suggested that ERAP1 coding SNPs can influence expression levels^47^. While it is not known if this applies to specific ERAP1 allotypes, it is not unlikely and should be considered when interpreting functional results, given that enzyme concentration is a major factor in determining enzyme kinetics. Finally, it is currently unknown how interactions between ERAP1 and other proteins in the ER, such as the chaperone ERp44^48^ modify its enzymatic activity. Regardless of these limitations, a robust understanding of the fundamental properties of ERAP1 as such is an obligatory step to build a more detailed framework of its functions in the more complex cellular setting. In this context our findings highlight that interpretations of ERAP1 functions in cellular or *in vivo* systems should be performed with caution, acknowledging the lack of knowledge of key parameters, such as substrate concentration.

In summary, we present evidence of key mechanistic differences that underlie the functional spectrum of the autoimmunity-protective ERAP1 allotype 10, highlighting the important role of two allotype-unique SNPs at positions 349 and 725. Allotype 10, appears to constitute a unique functional ERAP1 allotype, optimized for lower activity against shorter substrates, induced by tweaking of specific kinetic and thermodynamic parameters that likely control rate-limiting steps involved in the conformational equilibrium and dynamics of the enzyme. ERAP1 allotypic variation is emerging to be a key contributor to the variability of adaptive immune responses in close synergy with MHC-I polymorphic variation. Future studies will be necessary to fully understand the impact and necessity of this polymorphic variability and how it can influence therapeutic applications that focus on modulating antigen presentation^9^.

## Author Contributions

G.G. constructed the ERAP1 variants with help from M.T., generated recombinant proteins with help from A.M., designed and performed enzymatic assays and interpreted data. A.P. helped design the study, and interpreted data. E.S. conceived the study, designed experiments, interpreted data and wrote the manuscript with input from all authors.

## Data availability

Data points used to generate the graphs presented in this paper are available upon request to the corresponding author.

